# Guanosine tetraphosphate (ppGpp) promotes tolerance to excess light in *Arabidopsis thaliana*

**DOI:** 10.1101/2025.11.06.686924

**Authors:** Shanna Romand, Chen Hu, Aurélie Crepin, Wojciech Nawrocki, Cécile Lecampion, Brigitte Ksas, Sylvie Citerne, Michel Terese, Matteo Sugliani, Milena Kulasek, Stefano Caffarri, Roberta Croce, Michel Havaux, Ben Field

## Abstract

Guanosine tetraphosphate (ppGpp) is a hyperphosphorylated nucleotide originally discovered in prokaryotes and found in the chloroplasts of plants and algae. In plants, ppGpp regulates photosynthetic activity, which is important in acclimation to environmental stresses such as nitrogen limitation. However, the full range of stresses involving ppGpp signalling is not yet known. Here, we investigated the role of ppGpp accumulation in the acclimation of plants to excess light. We found that the over-accumulation of ppGpp in transgenic lines that overexpress *RSH3* (OX:RSH3) increases tolerance to excess light. Although ppGpp leads to higher non-photochemical energy dissipation (NPQ) than in wild-type plants, we show that NPQ is not critical for the enhanced excess light tolerance. Rather, our results show that ppGpp accumulation leads to broad changes in plant physiology that prime plants to resist and subsequently recover from excess light exposure. ppGpp levels themselves also increase in response to excess light exposure, suggesting that ppGpp may play a physiological role in excess light acclimation. Our work adds to a growing body of evidence for the importance of ppGpp in plant stress acclimation.

## Introduction

Sunlight is essential for driving photosynthesis and growth in plants. However, light quality and intensity can change rapidly, necessitating rapid acclimation mechanisms to optimise growth rates and protect cellular components. Indeed, the absorption of excess energy by the photosynthetic machinery can promote the over-excitation of chlorophyll and the formation of singlet oxygen (^1^O_2_) and other reactive oxygen species (ROS), sources of oxidative stress that damage lipids and proteins in the thylakoid membranes (Lee and Kim 2024).

Plants possess both constitutive and inducible mechanisms for dissipating the excess energy absorbed by the photosynthetic machinery harmlessly in the form of heat. The main inducible mechanism for energy dissipation is known as non-photochemical quenching (NPQ) (Bassi and Dall’Osto 2021; Ruban and Wilson 2021). NPQ consists of several components, including the well-known qE component. qE is induced within seconds by excess-light triggered acidification of the thylakoid lumen, reaches its maximum after a few minutes, and relaxes rapidly in the dark. During qE, lumen acidification is sensed by the PsbS protein, which is then thought to alter the conformation of the light-harvesting antenna (trimeric LHCII and monomeric Lhcb) of photosystem II (PSII) to promote quenching. Lumen acidification also activates the lumen-localised violaxanthin de-epoxidase (VDE) to produce the xanthophyll zeaxanthin, which enhances NPQ.

The ppGpp signalling pathway is emerging as another major stress acclimation mechanism in the chloroplast. ppGpp (or guanosine 3’,5’-diphosphate) is synthesised from ATP and GDP/GTP in the chloroplast by nuclear-encoded enzymes of the RelA/SpoT (RSH) homolog family (Boniecka et al. 2017; Mehrez et al. 2023). Arabidopsis encodes four RSH proteins, grouped into three subfamilies: RSH1, a ppGpp hydrolase that lacks synthase activity; RSH2 and RSH3, which function redundantly as bifunctional synthases/hydrolases; and the calcium-activated CRSH (Ca²⁺-dependent RSH) that is activated in the dark (Mehrez et al. 2023). We previously showed that RSH3 is directly controlled by the TOR kinase pathway, linking chloroplast ppGpp metabolism to central growth regulation (D’Alessandro et al. 2024).

In prokaryotes, ppGpp (and other 3’,5’-phosphorylated guanosine nucleotides) regulate bacterial growth and promote stress acclimation (Irving et al. 2021). ppGpp acts as a pleiotropic small-molecule effector that binds and modulates a broad range of target proteins involved in transcription, translation, and nucleotide metabolism-both competitively, at sites that normally accommodate GTP or GDP, and allosterically-to coordinate these responses. In plants, ppGpp regulates plant growth and development (Sugliani et al. 2016; Ono et al. 2021; Turkan et al. 2023), immunity (Abdelkefi et al. 2018; Qiu et al. 2023; Aballay et al. 2026), and responses to nitrogen deprivation (Maekawa et al. 2015; Honoki et al. 2018; Goto et al. 2022; Li et al. 2022a; Romand et al. 2022). Over-accumulation of ppGpp in the chloroplast down-regulates photosynthetic activity in both land plants (Maekawa et al. 2015; Sugliani et al. 2016; Honoki et al. 2018; Harchouni et al. 2022; Ito et al. 2022; Li et al. 2022a) and algae (Avilan et al. 2021). Consistent with its role in bacteria, ppGpp is thought to act in plants through the regulation of plastid transcription and translation (Mehrez et al. 2023), nucleotide metabolism (Nomura et al. 2014a, 2014b; Romand et al. 2022) and protein targeting (Zegarra et al. 2025). Recently, ppGpp was shown to play a physiological role in the acclimation of Arabidopsis to nitrogen starvation by inducing the remodelling of the photosynthetic electron transport chain to reduce photosynthetic activity and reduce oxidative stress (Romand et al. 2022). Intriguingly, ppGpp over-accumulation in Arabidopsis also promotes a strong NPQ (Honoki et al. 2018), suggesting a potential mechanism for reducing oxidative stress during nitrogen starvation.

The increased NPQ in response to ppGpp over-accumulation, along with its role in protecting plants from oxidative stress during nitrogen starvation, led us to ask whether ppGpp might contribute to excess light acclimation in Arabidopsis-a situation in which the photosynthetic machinery absorbs excess energy. In this study, we show that the accumulation of ppGpp in transgenic lines that overexpress *RSH3* (OX:RSH3) indeed increases tolerance to excess light. However, we show that NPQ is surprisingly not critical for this increased excess light tolerance. Rather, we find that ppGpp accumulation promotes excess light tolerance at multiple levels, effectively priming OX:RSH3 plants for exposure to excess light. Finally, we find that ppGpp increases in response to excess light in wild type plants, suggesting that ppGpp can play a direct role in the acclimation of plants to excess light stress.

## Results

### ppGpp accumulation improves excess light tolerance

ppGpp protects plants against photo-oxidative stress during nitrogen limitation (Romand et al. 2022). We therefore tested whether ppGpp over-accumulation alters the resistance of plants to excess light (EL), which also promotes photo-oxidative stress. EL was applied using a treatment that induces photobleaching in wild-type plants, combining photosynthetic photon flux density (PPFD) of 1500 μmol photons m⁻² s⁻¹ with a low air temperature (4–6 °C) that keeps leaf temperature close to growth temperature (see Materials and Methods). We exposed two independent RSH3 overexpression lines, OX:RSH3.1 and OX:RSH3.2 (Sugliani et al. 2016), which accumulated approximately 4- and 2.5-fold more ppGpp than the wild type under our conditions (Fig. S1). After 48 h of EL, the wild type showed extensive photobleaching, while both OX:RSH3 lines showed almost none (Fig. 1a,b). In agreement with this observation, chemiluminescence imaging of peroxidised lipids revealed high levels in the wild type but not in OX:RSH3 plants (Fig. 1c). Quantification of hydroxy-octadecatrienoic acids (HOTEs), the oxidation products of linolenic acid, the main fatty acid in Arabidopsis leaves, confirmed these results: after EL, HOTE levels in the wild type were around three times higher than in either OX:RSH3 line (Fig. 1d). These results show that ppGpp over-accumulation reduces oxidative damage under EL.

**Figure 1.**
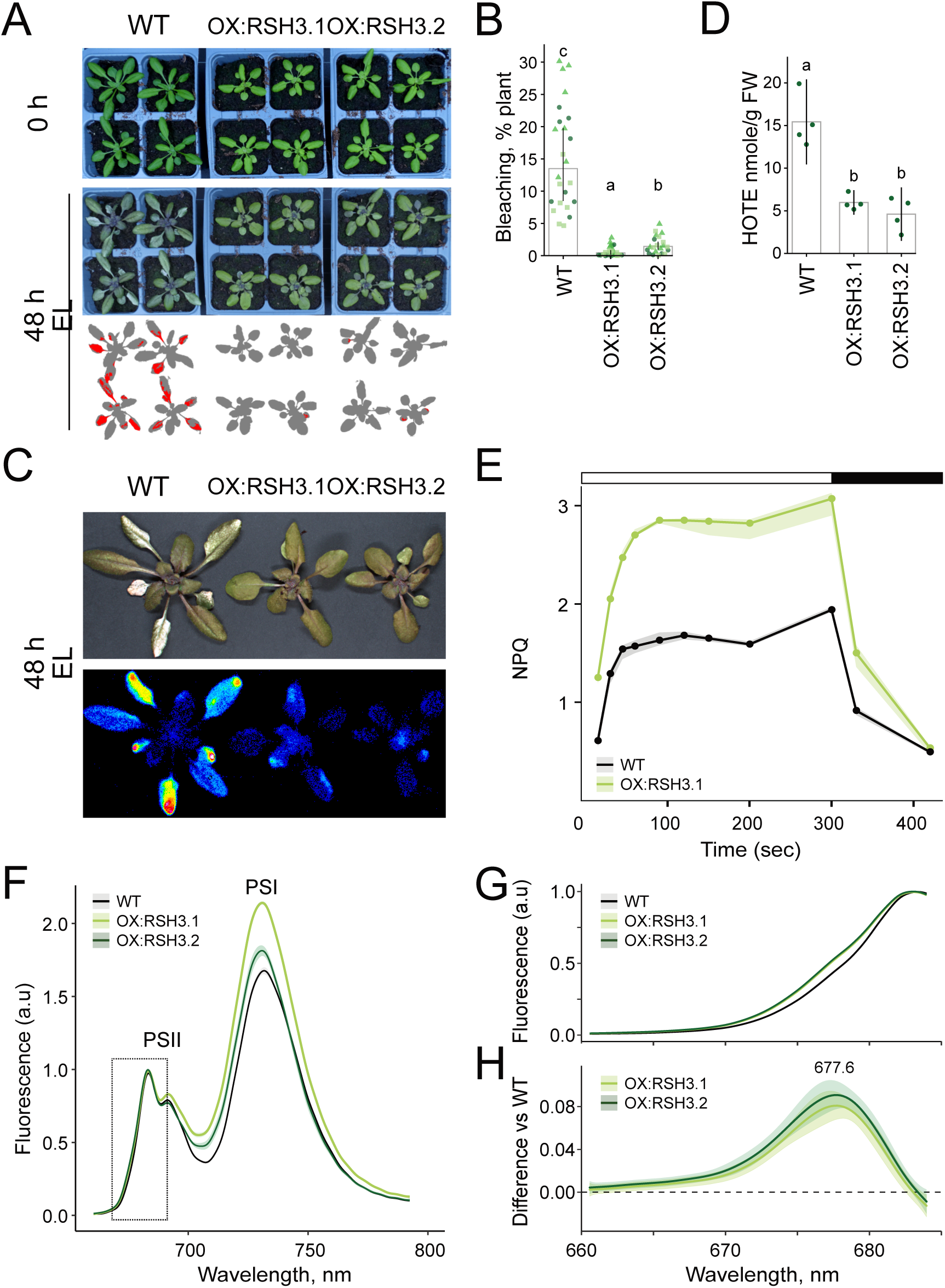
OX:RSH3 lines are tolerant to excess light and show increased NPQ and altered PSII antenna organisation. (A) Wild type and two independent OX:RSH3 lines before (0 h) and after 48 h exposure to excess light (EL; 1500 PPFD, air temperature 4–6 °C). The bottom row shows threshold detection of photobleached areas (red) on each plant (grey). (B) Quantification of photobleaching after 48 h EL. (C) Lipid peroxidation in plants after 48 h EL, visualised by chemiluminescence imaging. Top, colour photograph; bottom, chemiluminescence image of the same plants after 30 min dark adaptation, shown in false colour (black, no signal; blue to red, increasing signal). (D) Hydroxy-octadecatrienoic acid (HOTE) content after 48 h EL. (E) NPQ kinetics in 4-week-old plants during exposure to actinic light of 1200 PPFD (white bar) followed by a relaxation phase in the dark (black bar). Data in panels 1E, 2A and S5 come from the same experiment. (F) In vivo chlorophyll fluorescence emission spectra at 77 K, measured in state I. Peaks for PSII and PSI chlorophyll are indicated; spectra were normalised to the PSII peak at 685 nm. (G) Enlarged view of the PSII region (660– 685 nm). (H) Difference spectra across the PSII region relative to the wild type. Graphs show medians ± 95% confidence intervals, except (D), which shows means ± 95% confidence intervals. n = 24 plants from three experimental replicates, indicated by point shape (B); n = 4 biological replicates (D–H). Differences were tested by aligned-ranks ANOVA (B) or ANOVA with Tukey’s post-hoc test (D) (see Materials and Methods); letters indicate statistical groups. For (F–H), contrasts with the wild type were assessed using difference spectra with simultaneous confidence intervals (see Materials and Methods). PPFD, photosynthetic photon flux density (µmol photons m⁻² s⁻¹).

### Alterations in the photosynthetic machinery in OX:RSH3 plants

Previously, another RSH3 overexpression line was shown to induce high levels of NPQ (Honoki et al. 2018). Higher NPQ levels could potentially explain the reduced oxidative damage and general excess light tolerance we observe in OX:RSH3 plants. We therefore measured NPQ in OX:RSH3.1: it showed a higher NPQ capacity when grown under control conditions compared to the wild-type line (Fig. 1e). These results demonstrate that elevated ppGpp levels robustly induce a downstream increase in photoprotective capacity.

In addition to NPQ, RSH3 overexpresser plants are known to show changes in photosystem architecture, with notably a reduced quantity of PSII core relative to PSII peripheral antenna, and reduced PSII functionality with a low Fv/Fm ratio due to increased basal fluorescence (F0) (Sugliani et al., 2016). We further investigated these changes to understand whether they could be involved in excess light tolerance. Immunoblotting and protein staining confirmed previous observations that the two OX:RSH3 lines have a low PSII core to peripheral antenna ratio (Fig. S2, CP43 versus Lhcb1,2). The effects of ppGpp over-accumulation on the photosystem I (PSI) core to peripheral antenna ratio were less clear (PsaA versus Lhca1), as it was lower in OX:RSH3.1 and similar to the wild-type control in OX:RSH3.2. OX:RSH3.1 has higher ppGpp levels than OX:RSH3.2 (Fig. S1), and biochemical data also previously suggested lower amounts of PSI core complex subunits in OX:RSH3.1 (Sugliani et al. 2016). Reduced quantities of subunits for cytochrome b6f (Cyt f) and ATP synthase (AtpC) were also observed in both RSH3 over-expression lines. A marked difference compared to previous studies on RSH3 overexpression lines is that Rubisco large subunit (RBCL) accumulation was similar to that in the wild type control. This is very likely due to differences in protein normalisation, which was performed against total chlorophyll levels here compared to against total protein content previously. Finally, immunoblotting showed that levels of PsbS, a key protein in NPQ, were higher in both RSH3 overexpression lines.

Photosystem parameters were then estimated using electrochromic shift measurements in the presence of inhibitors. We noted an increase in PSII photochemical rate in the high ppGpp lines compared to the wild type, although this was significant only for OX:RSH3.2 (Fig. S3). An increased PSII photochemical rate is consistent with the presence of more peripheral antennae per PSII core complex, as also shown by immunoblotting here (Fig. S2) and previously (Sugliani et al. 2016). The PSI rate was significantly faster in both lines, suggesting likewise that there are more antenna per PSI core complex. The effect of ppGpp accumulation did not cause a striking change in the relative quantity of PSI (the PSI/(PSI+PSII) ratio).

Finally, we measured in vivo chlorophyll fluorescence emission at 77 K in plants in state 1. Analysis of the fluorescence spectra revealed a significant increase in the shoulder on the short-wavelength side of the PSII emission peak in RSH3 overexpressers compared to the wild type (Fig. 1f–h; global test, P < 0.00025). The maximal difference was observed between 677 and 678 nm (95% confidence interval). LHCII trimers emit at lower wavelengths than the PSII core, indicating that in the overexpressers, part of the peripheral antenna is less efficiently coupled to the PSII core. Taken together with the electrochromic shift measurements and immunoblotting, these results suggest that RSH3 overexpressers not only have an increased antenna-to-PSII core ratio, but also retain a fraction of peripheral antenna complexes that are more loosely connected to the core.

### OX:RSH3 tolerance to sustained excess light is largely independent of NPQ

The increased NPQ in OX:RSH3 lines appeared to be a plausible mechanism for their tolerance to sustained excess light. The rapid induction and dark relaxation kinetics suggest that the higher NPQ is due to an upregulation of the qE component of NPQ, which is dependent on PsbS and VDE. We therefore crossed OX:RSH3.1 with the *npq4* mutant, which lacks PsbS, and the *npq1* mutant, which lacks VDE, to obtain OX:RSH3.1 *npq4* and OX:RSH3.1 *npq1* double mutants, and used these to test whether NPQ was required for excess light tolerance.

We first confirmed that the *npq1* and *npq4* mutations in the OX:RSH3.1 genetic background effectively suppressed the high NPQ phenotype (Fig. 2a). We also verified that the double mutants maintained elevated ppGpp levels (Fig. S1). ppGpp levels in OX:RSH3.1 *npq4* were similar to OX:RSH3.1, while levels in OX:RSH3.1 *npq1* were moderately lower than in OX:RSH3.1 but similar to OX:RSH3.2. Interestingly, both double mutant lines were larger than the parental OX:RSH3.1, and OX:RSH3.1 *npq1* was also greener (Fig. S4a). These differences were more pronounced under short-day conditions at the same light intensity (Fig. S4b). The larger size of OX:RSH3.1 *npq4* cannot be explained by a change in ppGpp levels, indicating that loss of NPQ components can partially relieve the growth restriction of OX:RSH3.1. In OX:RSH3.1 *npq1*, the somewhat lower ppGpp levels may also contribute. We did not analyse this growth phenotype further here, and instead focused on whether the double mutants retained the excess light tolerance of OX:RSH3.1.

**Figure 2.**
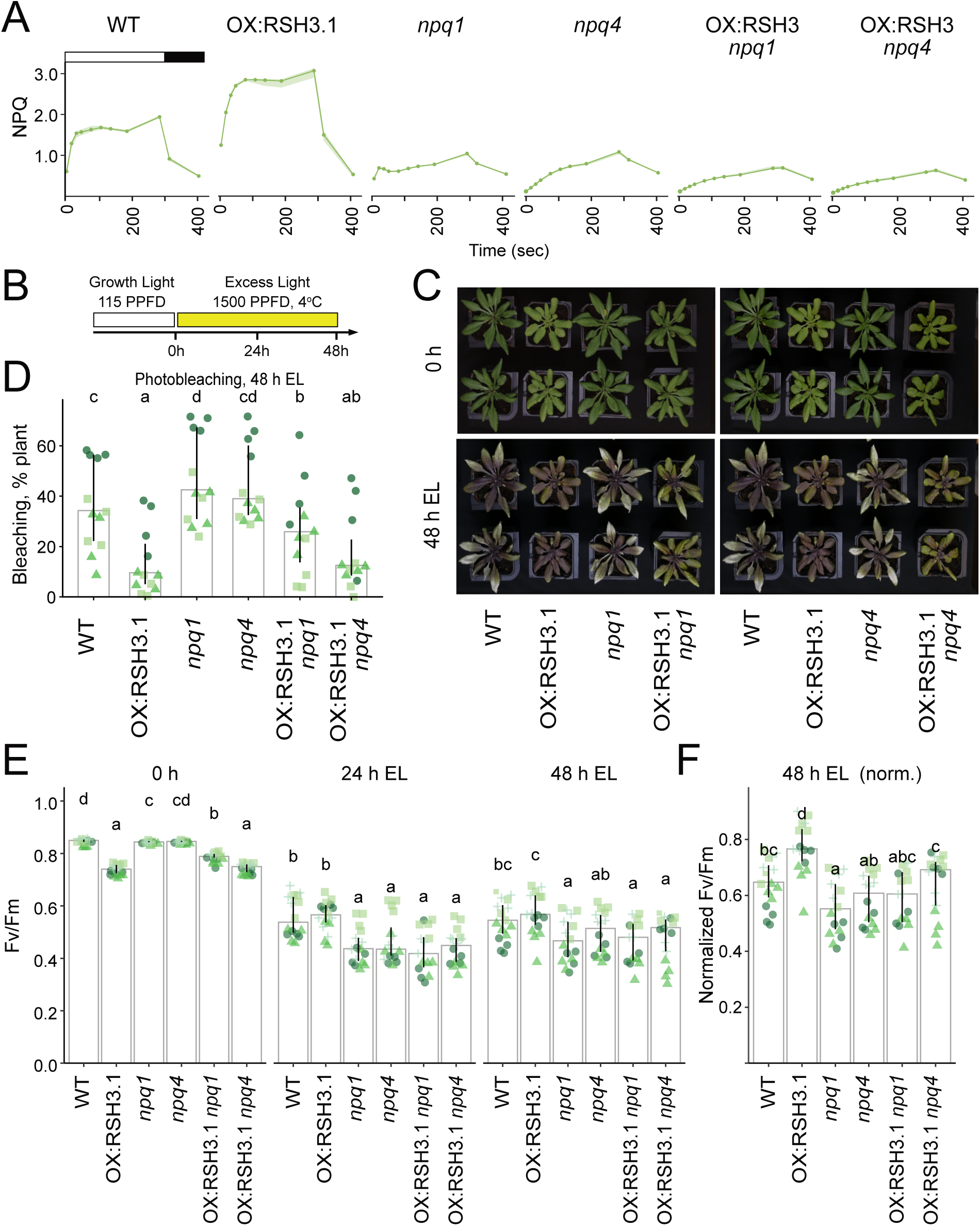
PsbS and VDE are not required for the tolerance of OX:RSH3 to excess light. (A) NPQ kinetics in 4-week-old plants of the indicated genotypes during exposure to actinic light of 1200 PPFD (white bar) followed by a relaxation phase in the dark (black bar). Curves show medians ± 95% confidence intervals, n = 4 plants. Data in panels 1E, 2A and S5 come from the same experiment. (B) Experimental protocol for 48 h EL exposure. Plants grown for 4 weeks under long-day (LD) conditions at 115 PPFD were exposed to excess light (EL; 1500 PPFD, air temperature 4–6 °C) for 48 h. (C) Representative plants before (0 h) and after 48 h EL. For clarity, the *npq1* and *npq4* series are shown in separate panels. All genotypes were grown and treated together, so the same wild-type and OX:RSH3.1 control plants appear in both panels. (D) Quantification of photobleaching after 48 h EL. (E) Maximum PSII yield (Fv/Fm) before EL and after 24 h and 48 h EL. (F) Fv/Fm after 48 h EL normalised to the 0 h value of each genotype. Data in (C–F) come from four independent experiments performed in three different growth facilities, with similar results. Graphs (D-F) show medians ± 95% confidence intervals for 3–4 experimental replicates (circles, triangles, squares and crosses). Differences were tested using aligned-ranks ANOVA with post-hoc pairwise comparisons (see Materials and Methods); letters indicate statistical groups between genotypes at each timepoint. PPFD, photosynthetic photon flux density (µmol photons m⁻² s⁻¹).

Wild-type plants as well as the *npq1* and *npq4* mutant lines showed extensive photobleaching after 48 h (Fig. 2b-d). OX:RSH3.1 again showed very little photobleaching compared to the wild type. The double mutant line OX:RSH3.1 *npq4* showed a similar level of photobleaching to OX:RSH3.1, and significantly less than the wild type or *npq4*. OX:RSH3.1 *npq1* showed more photobleaching than OX:RSH3.1, possibly due to the loss of zeaxanthin or its slightly lower ppGpp levels (Fig. S1), but photobleaching remained significantly lower than in the wild type and *npq1*. Therefore, removal of PsbS or VDE, and with it the high NPQ of OX:RSH3.1 (Fig. 2a), has a limited effect on the tolerance of OX:RSH3.1 to sustained excess light.

### ppGpp improves relative photosynthetic performance during prolonged excess light

Photobleaching is positively correlated with PSII photoinhibition, which occurs when the rate of oxidative damage to PsbA (a central protein of the PSII reaction centre) exceeds the rate of repair (Pinnola and Bassi 2018; Nawrocki et al. 2021). We therefore tested the susceptibility of RSH3 overexpressers to photoinhibition.

In control conditions, OX:RSH3 plants were small and pale, and showed a lower Fv/Fm ratio than the control (Fig. 2e), as previously observed (Sugliani et al., 2016). As discussed above, the low Fv/Fm in OX:RSH3 is a result of a high basal fluorescence (F0) caused by the disconnection of a fraction of LHCII from the PSII core, and is unlikely to reflect the direct impairment of PSII core function itself. The OX:RSH3 *npq4* lines showed a similar Fv/Fm ratio, while OX:RSH3 *npq1* showed a ratio between OX:RSH3 and the wild type.

After 24h excess light exposure, all lines showed a significant decrease in Fv/Fm as a result of photoinhibition. The OX:RSH3.1 *npq1* and OX:RSH3.1 *npq4* lines showed a similar response to *npq1* and *npq4*, with significantly lower Fv/Fm ratios than OX:RSH3.1 or the wild type (Fig. 2e). Absolute Fv/Fm ratios after 48 h were similar to those at 24 h. To compare the relative effects between lines we also calculated the Fv/Fm normalized to Fv/Fm before excess light treatment (0h). This showed that OX:RSH3.1 underwent a relatively lower level of photoinhibition than the wild type, while the double mutants were no longer distinguishable from the corresponding *npq* single mutants (Fig. 2f). Despite this photoinhibition, both double mutants still showed much less photobleaching than the wild type or the *npq* single mutants (Fig. 2d). Together, these results suggest that ppGpp accumulation improves relative photosynthetic performance under prolonged excess light, and that its protection against photobleaching does not require qE.

We also obtained NPQ kinetics after excess light stress exposure (Fig. S5). NPQ remained higher in OX:RSH3 than in the wild type even after excess light exposure. For *npq1*, *npq4*, OX:RSH3.1 *npq1* and OX:RSH3.1 *npq4* the kinetics were relatively similar to those obtained under control conditions, with a low maximum NPQ. Interestingly, for all lines we found that NPQ capacity decreased after exposure to excess light. This phenomenon likely reflects competition from photoinhibition-related qI quenching (Nawrocki et al. 2021), together with reduced PSII activity, which reduces lumen acidification and the triggering of qE.

### qE is important for the initial photoinhibition response of OX:RSH3 plants

Photobleaching and photoinhibition after 24-48h is the result of short and long-term processes, both of which may be influenced by ppGpp. To gain insights into the initial sensitivity of OX:RSH3 plants to excess light we therefore quantified photoinhibition in short time-course experiments on detached leaves (Fig. 3a,b).

**Figure 3.**
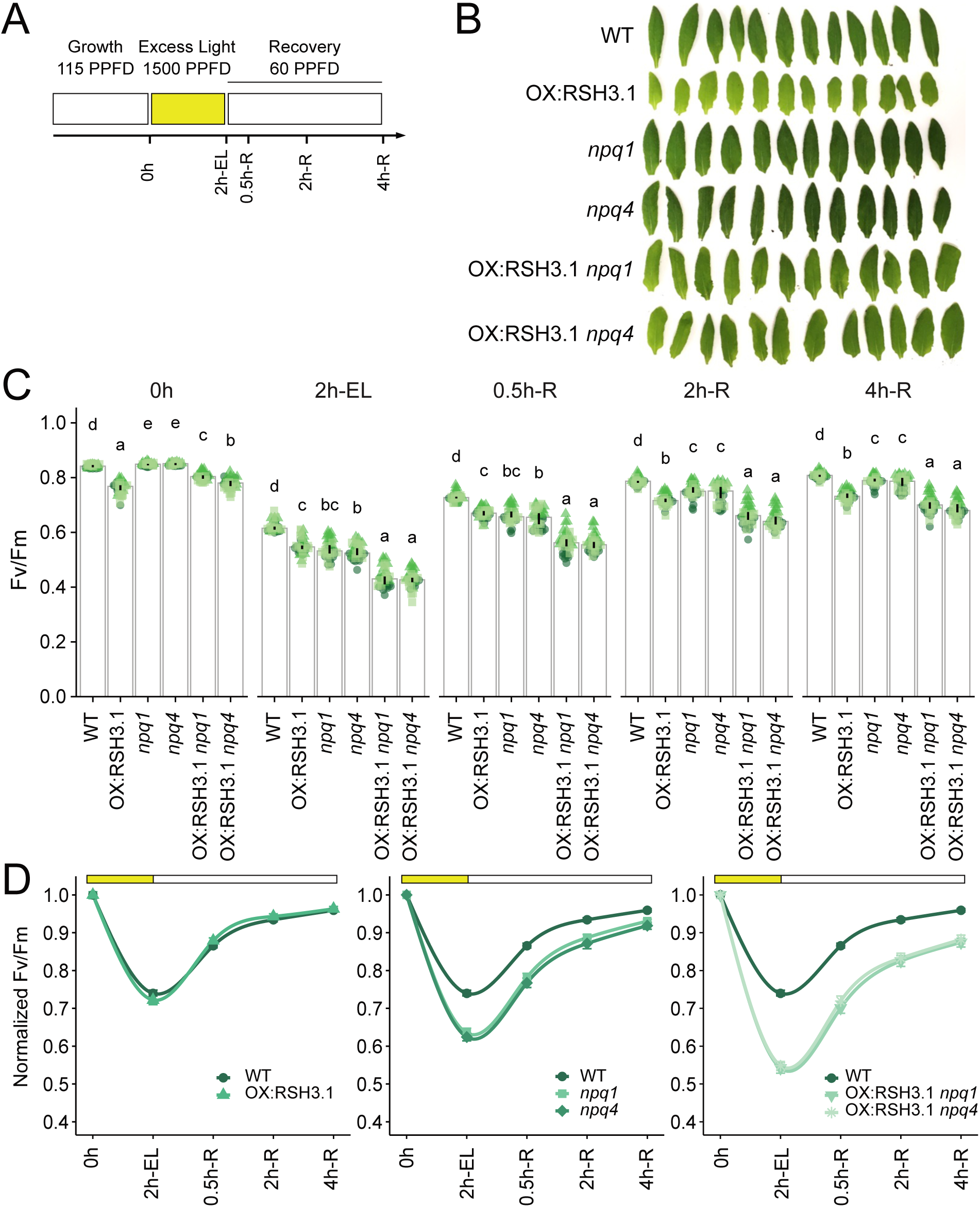
Removal of PsbS or VDE increases photoinhibition sensitivity in OX:RSH3.1 plants. (A) Experimental protocol. Detached leaves of the indicated genotypes were exposed to excess light (EL; 1500 PPFD, air temperature 4–6 °C) for 2 h and then allowed to recover at 60 PPFD at 22 °C. Timepoints: 0h, before EL; 2h-EL, after 2 h EL; 0.5h-R, 2h-R and 4h-R, after 0.5, 2 and 4 h of recovery. (B) Representative detached leaves of each genotype before EL exposure. (C) Maximum PSII yield (Fv/Fm). Graphs show medians ± 95% confidence intervals for three experimental replicates (circles, triangles and squares), each with 12 detached leaves per genotype. Differences were tested using aligned-ranks ANOVA and post-hoc pairwise comparisons (see Materials and Methods); letters indicate statistical groups between genotypes at each timepoint. (D) Fv/Fm normalised to the mean value before EL exposure (0h = 1). Symbols show means ± 95% confidence intervals and lines are smoothed fits shown as a visual guide. The same wild-type data are shown in each panel to allow comparison with each genotype. PPFD, photosynthetic photon flux density (µmol photons m⁻² s⁻¹).

Before EL exposure, both double mutants showed a higher Fv/Fm than OX:RSH3.1, with OX:RSH3.1 *npq1* closer to the wild type (Fig. 3c). This was associated with a lower F0 in the double mutants versus OX:RSH3.1, rather than a higher Fm (Fig. S6a). The difference in Fv/Fm was less clear for OX:RSH3.1 *npq4* in whole-plant measurements (Fig. 2e), which were more variable. After 2 h exposure to excess light, strong photoinhibition developed in all lines, and strikingly, Fv/Fm in OX:RSH3.1 *npq1* and OX:RSH3.1 *npq4* fell significantly below that of either parental line (Fig. 3c). Plotting the normalised Fv/Fm showed the same pattern. The relative responses of OX:RSH3.1 and the wild type to 2 h excess light were almost identical, dropping to 74% and 72% of starting Fv/Fm respectively, while *npq1* and *npq4* dropped further (68% and 64%) and the two double mutants dropped most (56% and 57%) (Fig. 3d, Fig. S6b). The same trend between the lines was also observed during recovery (Fig. 3c,d). Using F0 and Fm values normalised to T0 for each genotype, the decrease in Fv/Fm was mainly associated with a decline in Fm, with smaller changes in F0 (Fig. S6b). Interestingly, relative to its starting value, OX:RSH3.1 maintained a higher Fm than the wild type after excess light that also recovered faster, but its higher initial F0 meant that this was not reflected in the normalised Fv/Fm. In contrast, Fm declined further and recovered more slowly in both the *npq* single and double mutants (Fig. S6b). Overall, these results show that OX:RSH3 plants are at least as resistant to short-term photoinhibition as the wild type. However, the response of the *npq* double mutants indicates that the qE component of NPQ may play a more important role in protecting OX:RSH3 plants against short-term photoinhibition than in plants with wild-type ppGpp levels (Fig. 3c,d). This short-term qE dependence contrasts with the situation after 48 h, when the double mutants were as photoinhibited as the *npq* single mutants (Fig. 2f) yet remained largely protected from photobleaching (Fig. 2d). Together, these results indicate that ppGpp protects against oxidative damage downstream of PSII photoinhibition, independently of qE.

### Tolerance to excess light is not a general feature of low chlorophyll mutants

Another feature of OX:RSH3 plants that could limit photo-oxidative damage is their low chlorophyll content, which may reduce light absorption. We therefore asked whether excess light tolerance is a general feature of plants with low chlorophyll content like OX:RSH3.

To explore this possibility we tested two mutants with low chlorophyll, *chlorina 1* (*ch1*) and the *GOLDEN-LIKE* (*GLK*) double mutant *glk1,2*. *ch1* lacks chlorophyll b due to the presence of a mutation in the chlorophyllide a oxygenase gene (*CAO*) (Espineda et al., 1999). This leads to a lack of the majority of the LHCII proteins that require chlorophyll b for assembly (Reinsberg et al. 2001), resulting in the accumulation of PSII core without peripheral antenna (Havaux et al. 2007). *glk1,2* on the other hand, lacks the GLK1 and GLK2 transcription factors that regulate the expression of nuclear-encoded genes for photosynthesis and chlorophyll biosynthesis (Waters et al. 2009; Li et al. 2022b), has reduced chlorophyll content, and displays wild-type-like photosynthetic parameters (Liu et al. 2018). Both lines showed strong photobleaching and greater photoinhibition than the wild type or OX:RSH3.1 under EL (Figs S7, S8). EL sensitivity has not previously been reported for *glk1,2*. Low chlorophyll content alone therefore does not explain the EL tolerance of OX:RSH3 plants.

### ppGpp is involved in priming plants for excess light acclimation

To identify additional mechanisms that could explain ppGpp-induced excess light tolerance, we searched the published list of genes differentially expressed in 12-day-old OX:RSH3.1 seedlings grown under non-stress conditions (Abdelkefi et al. 2018) for genes associated with excess light or oxidative stress responses. These plants showed increased accumulation of transcripts for *EARLY LIGHT INDUCIBLE PROTEIN* (*ELIP1* and *ELIP2*), *HYDROPEROXIDE LYASE 1* (*HPL1*), *COPPER/ZINC SUPEROXIDE DISMUTASE 2* (*CSD2*), *STROMAL ASCORBATE PEROXIDASE* (*SAPX*) and methionine sulfoxide reductases (*MSRB6* and *MSRB7*). This suggests that even in the absence of stress, OX:RSH3.1 plants may be constitutively primed for tolerance to excess light exposure. To test whether these elevated transcript levels were reflected at the protein level, we tested ELIP2 protein levels in mature plants by immunoblotting. We observed significantly higher levels of ELIP2 in OX:RSH3.1 plants after a short exposure to excess light (Fig. 4a,b).

**Figure 4.**
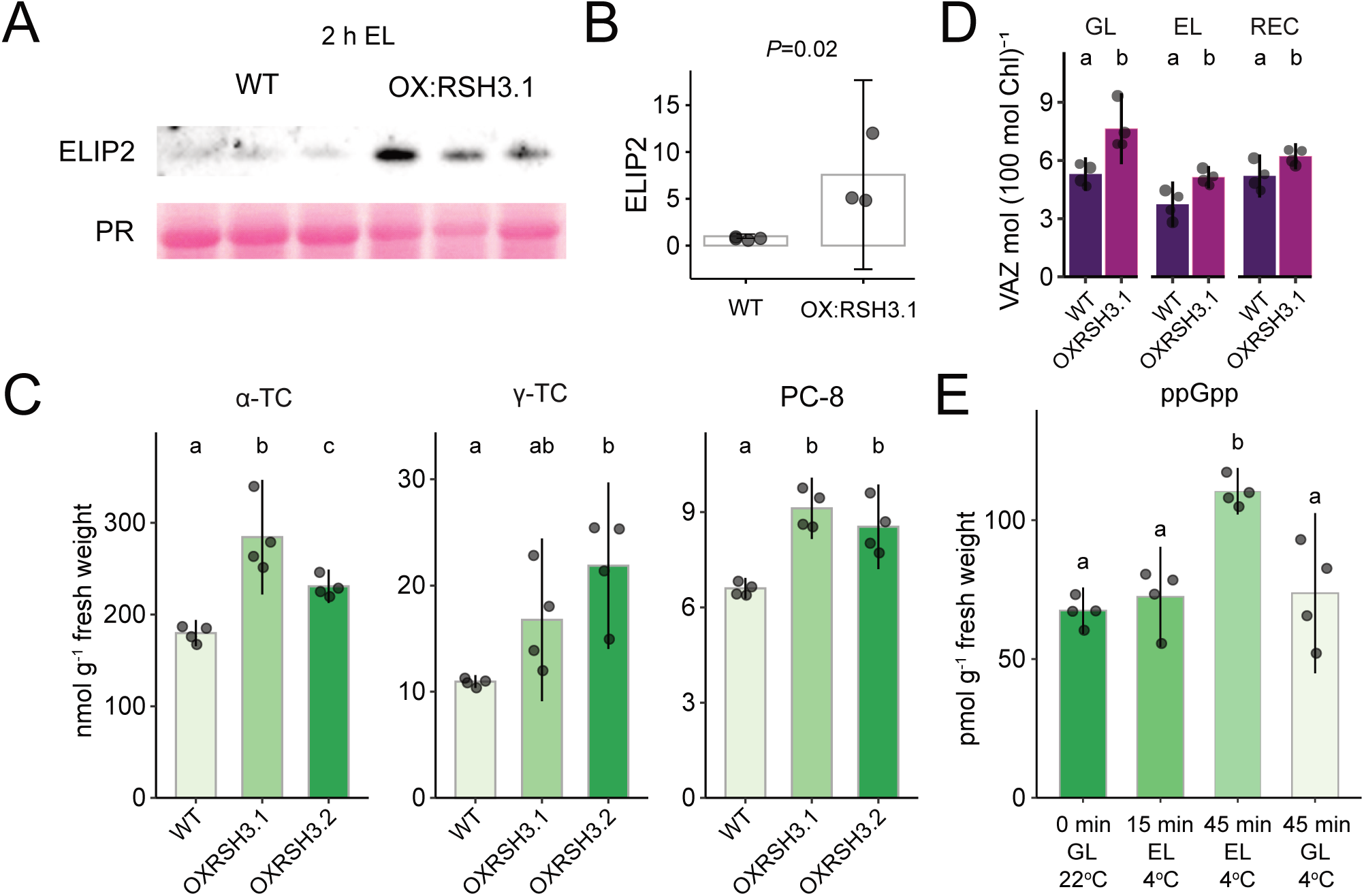
ppGpp primes plants for excess light acclimation. (A) Immunoblot showing the abundance of ELIP2 in 20 µg total protein extracted from 4-week old plants exposed to 2 h excess light (EL) stress (1500 μmol photons m⁻² s⁻¹ at 4–6 °C). Total proteins were revealed using Ponceau red stain (PR). (B) Quantification of ELIP2 relative protein abundance from immunoblots, 3-4 biological replicates. (C) Antioxidant content in 4-week old plants grown under standard growth conditions. α-TC, α-tocopherol; γ-TC, γ-tocopherol and PC-8, plastochromanol-8. (D) Combined amount of violaxanthin, antheraxanthin and zeaxanthin (VAZ) under growth light (GL), after 2 h EL exposure (EL) and 1 h of recovery under standard lighting (REC). Values normalized to 100 total chlorophyll (a+b). (E) ppGpp content in plants grown under standard conditions with growth light (GL) for 4 weeks and then exposed to 1500 μmol of photons m^-2^s^-1^ at 4–6 °C (EL) for the indicated times. Graphs show mean ± 95% confidence interval, n = 3-4 biological replicates. Differences were tested using ANOVA with Tukey’s post-hoc test (see Materials and Methods). Letters indicate statistical groups.

We next examined whether antioxidant pools were also affected. Tocopherols and plastochromanol-8 are important lipophilic antioxidants capable of quenching singlet oxygen (Havaux 2020). α-Tocopherol, the most abundant antioxidant, accumulated to significantly higher levels in OX:RSH3.1 than in the wild type under standard non-stress growth conditions, with intermediate levels in OX:RSH3.2 (Fig. 4c). γ-Tocopherol and plastochromanol-8 also showed a significant increase in both OX:RSH3 lines compared to the wild type.

We then turned to the role of pigments. Under growth light, OX:RSH3.1 plants had a significantly larger VAZ (violaxanthin, antheraxanthin and zeaxanthin) pool than the wild type (Fig. 4d), with higher levels of antheraxanthin, an intermediate in zeaxanthin synthesis (Fig. S9). Zeaxanthin, which usually accumulates only under excess light, was even detected in one of the OX:RSH3.1 replicates. The VAZ pool remained larger in OX:RSH3.1 than in the wild type after a short EL treatment and during recovery (Fig. 4d). During recovery, zeaxanthin and antheraxanthin levels stayed higher, leading to a higher de-epoxidation state (DEPS) (Fig. S9). Thus, ppGpp over-accumulation increases the xanthophyll cycle pool both before, during and after EL exposure.

### ppGpp levels increase rapidly in response to excess light

Finally, we asked whether ppGpp levels themselves respond to excess light. In wild-type plants, ppGpp levels were unchanged after 15 min of EL, but increased significantly after 45 minutes. They remained unchanged in plants kept under growth light at the same low air temperature (Fig. 4e). The dynamics beyond this point were not determined.

## Discussion

Here, we show that ppGpp accumulation promotes tolerance to excess light in Arabidopsis. We first extended previous work characterizing the photosynthetic phenotype of ppGpp-overaccumulating plants (Sugliani et al. 2016; Honoki et al. 2018), providing additional data on photosynthetic complex composition, and confirming the elevated NPQ capacity of these plants (Fig. 1). We then found that while the qE component of NPQ is particularly important for moderating sensitivity to photoinhibition in ppGpp over-accumulators, it is not essential for resistance to photobleaching (Figs 2, 3). Although OX:RSH3 plants have reduced chlorophyll content, low chlorophyll alone did not confer excess light tolerance, as two other pale lines with different photosynthetic defects were highly sensitive (Figs S7, S8). Rather, ppGpp over-accumulators showed evidence of constitutive priming for excess light resistance with increased expression of genes responsive to excess light or oxidative stress, elevated ELIP2 levels, larger antioxidant pools and an enlarged xanthophyll cycle pool (Fig. 4). ppGpp therefore appears to promote a broad spectrum of responses, including NPQ, that together confer resilience against excess light. Interestingly, ppGpp levels themselves also rise in response to excess light exposure (Fig. 4e). Together with the protective effect of ppGpp over-accumulation, this suggests that ppGpp may play a physiological role in excess light acclimation. Determining the excess light sensitivity of ppGpp-deficient lines (Inazu et al. 2024) will be an important next step, as will characterising the longer-term kinetics of the ppGpp response. This adds to the growing list of roles for ppGpp in plant physiology (Mehrez et al. 2023), and underscores its importance in plant stress acclimation.

How is ppGpp able to induce constitutive priming? The principal known mode of action for ppGpp signalling in vivo involves repression of chloroplast transcription and translation (Mehrez et al. 2023), that is, plastid gene expression. Repression of plastid gene expression is also a well-known trigger for retrograde signalling to the nucleus (Liebers et al. 2022). However, ppGpp-mediated repression of plastid gene expression occurs in mature chloroplasts and in response to stress, so any resulting signal might not fit neatly into the classical distinction between biogenic and operational retrograde signalling. Nevertheless, such signalling provides a plausible route through which ppGpp could activate the transcription of excess light and oxidative-stress responsive genes. Beyond this, in vitro studies have also implicated chloroplast nucleotide metabolism and protein trafficking as additional targets of ppGpp (Nomura et al. 2014a, 2014b; Zegarra et al. 2025), suggesting that multiple pathways may influence the priming response.

ppGpp accumulation also induces changes in the architecture of the photosynthetic machinery, such as the relative enlargement of the PSII peripheral antenna system. Counterintuitively, the presence of disconnected antenna would be expected to increase sensitivity by enhancing singlet oxygen production, yet OX:RSH3 plants are more resistant to excess light. This suggests that additional protective mechanisms must outweigh this potential risk. In particular, the LHCII antenna-although partially disconnected from PSII in OX:RSH3 and therefore expected to be vulnerable -appears to be well protected, at least in part through mechanisms associated with the VAZ xanthophyll cycle and PsbS, which we show are enhanced by ppGpp accumulation (Figs S2, S9). Increased ppGpp levels might also be expected to disrupt the repair cycle of the PSII RC protein D1, which is critical during photoinhibition and recovery. Surprisingly, however, OX:RSH3 plants show similar or slightly faster photoinhibition recovery than the wild type, suggesting that ppGpp does not have a major impact on the repair cycle (Fig. 3d, Fig. S6). Nevertheless, removal of VDE or PsbS (in *npq1* and *npq4,* respectively) leads to stronger photoinhibition and slower recovery in OX:RSH3 than in either mutant alone (Fig. 3d). This indicates an interaction between ppGpp signalling and qE, which we speculate may relate to the enlarged PSII peripheral antenna system in ppGpp-accumulating plants. In conclusion, while the qE component of NPQ is important for the initial development of photoinhibition in OX:RSH3 plants, our results reveal that it is only one of several excess light acclimation processes triggered by ppGpp accumulation. Together, these processes contribute to the enhanced excess light resistance observed in ppGpp-accumulating plants.

While these mechanisms enhance excess light resistance in OX:RSH3 plants, they also appear to come at a cost to growth. Over-accumulation of ppGpp often leads to reduced plant growth, at least under standard nutrient-replete growth conditions (Sugliani et al. 2016; Goto et al. 2022; Li et al. 2022a). Interestingly, loss of PsbS or VDE partially relieved the growth restriction of OX:RSH3.1 (Fig. S4), and both double mutants showed a higher Fv/Fm than OX:RSH3.1 under control conditions (Fig. 3c). This partial reversion was stronger in OX:RSH3.1 *npq1*, possibly due to its lower ppGpp levels. However, it also occurred in OX:RSH3.1 *npq4*, where ppGpp levels were unchanged (Fig. S1). This suggests that the enhanced NPQ capacity of ppGpp over-accumulating plants may limit growth by dissipating energy that could otherwise be used for photosynthesis. Consistent with this view, under low light, plants lacking PsbS accumulate more biomass than wild-type plants (Khuong et al. 2019), while constitutive zeaxanthin accumulation in the *npq2* mutant is associated with reduced growth (Dall’Osto et al. 2005).

## Acknowledgements

We thank Muriel Reissolet and Patrice Crete for plant care, Chloé Maurin and Theo Mege for assistance with immunoblotting, and Julia Bartoli and Emmanuelle Bouveret for supplying isotope labeled ppGpp. Nucleotide measurements were performed on the IJPB Plant Observatory technological platform with support from Saclay Plant Sciences-SPS (ANR-17-EUR-0007) and at Nicolaus Copernicus University Toruń. Work was funded by the Agence Nationale de la Recherche (ANR-17-CE13-0005, ANR-22-CE20-0033), and work at VU Amsterdam was supported by the Dutch Organization for Scientific research via a TOP grant.

## Material and Methods

### Plant material and growing conditions

All *Arabidopsis thaliana* lines used in this study are listed in Table 1. Seeds were sown and germinated on potting soil and the seedlings were transferred to individual pots 7 days after germination. The plants were then grown in a growth chamber with a photoperiod of 16h light (at 22°C) for 8h darkness (at 19.5°C) with LED illumination at a PPFD of 115, the plants received two weekly waterings and a weekly application of Coïc-Lesaint fertilizer solution.

**Table 1:** Lines used in this study.

| **Lines** | **Designation** | **Source or reference** | **Additional information** |
|---|---|---|---|
| Col-0 | WT, Columbia, N1093 (NASC) | Nottingham Arabidopsis |  |
| OX:RSH3.1 | OX:RSH3.1 (H1) | (Sugliani et al. 2016) | RSH3 overexpression line (35S:RSH3-GFP), genetic background Col-0. |
| OX:RSH3.2 | OX:RSH3.2 (G8) | (Sugliani et al. 2016) | RSH3 overexpression line (35S:RSH3-GFP), genetic background Col-0 |
| *npq1* | *npq1-2* | (Niyogi et al. 1998) | Genetic background Col-0 |
| *npq4* | *npq4-1* | (Li et al. 2000) | Genetic background Col-0 |
| OX:RSH3.1 *npq1* | OX:RSH3.1 *npq1-2* | This study |  |
| OX:RSH3.1 *npq4* | OX:RSH3.1 *npq4-1* | This study |  |
| *ch1* | *ch1-1* | (Espineda et al. 1999) |  |
| *glk1,2* | *glk1,2* | (Fitter et al. 2002) | Seeds were provided by Jane Langdale |

To create the double mutants OX:RSH3.1 was crossed with *npq1-2*, *npq4-1* and *ch1-1*. In the F2 generation, plants carrying the RSH3 overexpression transgene were identified by BASTA (glufosinate ammonium) resistance and the characteristic pale green phenotype of OX:RSH3 plants. *npq1* and *npq4* homozygous mutants were identified by the absence of NPQ induction using chlorophyll fluorescence imaging, and *ch1* homozygous mutants by their chlorophyll b-deficient phenotype. Lines homozygous for the transgene were then selected by the absence of BASTA-sensitive segregants among their progeny. Over-accumulation of ppGpp in the *npq1* and *npq4* double mutants was confirmed by direct quantification (Fig. S1). In contrast, ppGpp levels in OX:RSH3.1 *ch1* were not significantly different from the wild type (Fig. S1), so this line was not used to assess the effect of ppGpp in a *ch1* background.

### Exposure to excess light

Plants or detached leaves were placed in a controlled environment culture chamber under LED lighting set at a continuous PPFD of 1500 μmol photons m^-2^ s^-1^ with the air temperature set to 4–6 °C. Under these conditions, leaf temperature remained at approximately 19–22 °C, close to the growth temperature: the low air temperature therefore compensates for leaf warming under strong illumination and avoids heat stress. For detached leaf experiments, the 5^th^ and 6^th^ rosette leaves were selected.

### Quantification of photobleaching

Photobleaching was quantified from images of plant trays in Fiji (Schindelin et al. 2012). Rosettes were segmented by colour thresholding, and the resulting mask was corrected manually to separate touching plants and to rejoin detached leaves, giving one region of interest (ROI) per rosette. Bleached tissue was segmented from the same image by colour thresholding. For each ROI, the total rosette area and the bleached area were measured, and photobleaching was expressed as the percentage of the rosette area that was bleached. Threshold values were adjusted for each experiment to account for differences in lighting and camera settings, but were kept identical for all genotypes within an experiment.

### Chemiluminescence imaging of lipid peroxidation

Spontaneous photon emission from peroxidised lipids was imaged after 48 h EL as described (Havaux and Ksas 2022). Plants were dark-adapted for 30 min to allow chlorophyll luminescence to decay, then imaged for 30 min with a Fusion Fx imager equipped with a cooled CCD camera (Vilber). Isolated hot pixels were removed, and images are shown in false colour.

### Quantification of HOTEs

Lipids were extracted from about 300 mg of leaves in methanol/chloroform and analyzed by HPLC–UV as described previously (Montillet et al. 2004; Shumbe et al. 2017).

### Chlorophyll fluorescence imaging

Plants were placed in the dark for 20 min and chlorophyll fluorescence was measured using the Fluorcam FC 800-O fluorometer (Photon system Instruments). The maximum quantum yield of PSII was calculated according to the formula: (Fm-Fo)/Fm. NPQ was calculated according to the formula NPQ = (Fm-Fm’)/Fm’.

### Measurement of the chlorophyll fluorescence emission spectrum at 77K

4-week-old plants were placed under far-red light for 60 min, to induce state I, then harvested and ground in liquid nitrogen. 50 μL of buffer (100% (w/v) glycerol, 10 mM Hepes -KOH, pH 7.5) was added to 30 mg of sample in a Pasteur pipette. Chlorophyll fluorescence spectra were obtained using a CARY Eclipse fluorescence spectrophotometer (Varian). The excitation light was set to 440 nm (maximum absorption wavelength of chlorophyll). The emission spectrum was measured by scanning between 600 and 800 nm. The spectra were normalised to the peak at 685 nm (max PSII emission).

### Immunoblotting

Protein extraction and immunoblotting were performed as described previously (Sugliani et al. 2016). Proteins were normalized on a total chlorophyll or total protein basis. Primary antibodies (supplied by Agrisera) were used against AtpC, CP43, Cyt f, ELIP2, Lhca1, PsaA and PsbS. ELIP2 was quantified after normalising ELIP2 signal to total protein (PR) for each sample.

### ECS measurements

The measurements were performed using a JTS-10 spectrophotometer as described (Hu et al. 2023), based on the method reported (Bailleul et al. 2010).

### Pigment quantification

Pigments were extracted by addition of acetone 90% 10 mM Tris HCl pH 7.5 to 6 mm leaf discs, followed by a grinding with glass beads for 10 sec at a speed of 6 m.s^-1^ in a bead beater (Bead Ruptor Elite, Omni International), and a 10 min centrifugation at maximum speed. The supernatant was passed through HPLC according to a protocol described previously (Campoli et al. 2009). Pigment amounts are expressed as mol per 100 mol total chlorophyll (a+b). Deepoxidation state (DEPS) was calculated as (Z+0.5A)/VAZ, where Z = zeaxanthin, A = antheraxanthin and V = violaxanthin.

### Quantification of antioxidants

Antioxidants (α- and γ-tocopherol, plastochromanol (PC)-8) were extracted from leaf tissue and quantified by reverse-phase HPLC as previously described (Ksas et al. 2015). In brief, leaf samples were ground in ethyl acetate. After centrifugation, the supernatant was filtered and evaporated on ice under a stream of N_2_. The residue was recovered in methanol/hexane (17/1) and analyzed by HPLC with fluorescence detection. Separation of tocopherols and PC was performed on a Phenomenex Kinetex column (2.6 µm, 100 × 4.6 mm, 100 Å) in the isocratic mode with methanol/hexane (17/1) as the solvent system and a flow rate of 0.8 ml min^−1^. Tocopherols and PC were detected by their fluorescence at 330 nm with excitation at 290 nm. PC standard was a kind gift from Dr. J. Kruk (Krakow, Poland). Tocopherol standards were purchased from Sigma.

### Guanosine tetraphosphate quantification

For the time-course experiment (Fig. 4e), nucleotides were extracted and quantified by HPLC– MS/MS using a stable isotope labelled ppGpp standard as described previously (Bartoli et al. 2020). The concentration of the in-house ¹³C-labelled ppGpp standard was calibrated against a commercial ppGpp standard.

For the comparison of genotypes (Fig. S1), nucleotides were extracted as described previously (Bartoli et al. 2020) and analysed by LC-MS/MS on a Nexera UHPLC coupled to an LCMS-8045 triple quadrupole mass spectrometer (Shimadzu). Separation was performed on an Ascentis Express C18 column (Supelco) at a flow rate of 0.3 mL min⁻¹. Solvent A was 8 mM hexylamine adjusted to pH 8.5 with acetic acid, and solvent B was acetonitrile. The gradient was as follows: 100% A at 0 min, decreasing linearly to 60% A at 10 min; a wash at 30% A at 12 min; a return to 100% A at 12.5 min; and re-equilibration at 100% A until 30 min. The electrospray ion source was operated in positive ion mode, and nucleotides were detected in multiple reaction monitoring (MRM) mode using the following transitions: GTP, m/z 524.0 → 152.0; ¹³C-GTP, 534.0 → 157.3; ppGpp, 603.8 → 152.1; ¹³C-ppGpp, 613.9 → 157.1. ppGpp and GTP were quantified relative to the ¹³C-labelled internal standards.

Because the two sets of measurements were performed on different instruments and with separate calibrations, absolute ppGpp levels are compared only within each experiment.

### Data analysis

Analyses were performed in R (R Development Core Team, 2020) using custom annotated R scripts. Command-line scripts were created to pre-process data obtained from the Fluorcam O-800, extract parameters, and calculate the photosynthetic indices. Graphs were produced using the ggplot2 package (Wickham 2009) with confidence intervals determined for normally distributed data using the Rmisc package (Hope 2013), and non-parametric data with bootstrap confidence interval estimation using the boot package (Davison and Hinkley 1997; Canty and Ripley 2021). Statistical analyses were performed on normally distributed data using analysis of variance (ANOVA) followed by Tukey’s post hoc test (Kassambara 2021). Non-parametric data were analysed using the Kruskal-Wallis test followed by Dunn’s post-hoc test (Kassambara 2021) for single-factor comparisons, or aligned-ranks ANOVA (Wobbrock et al. 2011) followed by post-hoc pairwise comparisons using emmeans (Lenth 2024) when experimental replicate was included as a random factor. 77K spectra (Fig. 1F-H) were analyzed in R using median quantile GAMs with line-specific smooths, and contrasts versus wild type were assessed by difference spectra with simultaneous confidence intervals and global supremum tests (Fasiolo et al. 2021). In figure panels 1B, 2D-F, 3C, S6A-B, S7C-E, and S8B the experimental replicates were included in the aligned-ranks ANOVA model as a random factor. The NPQ data shown in Figures 1E, 2A, and S5 were obtained from the same experiment; wild type and OX:RSH3.1 control data are therefore identical across these panels. All statistical tests included adjustments for multiple comparisons.

## Data availability

Raw and processed data for all figures are available in the Recherche Data Gouv repository under temporary DOI: https://entrepot.recherche.data.gouv.fr/dataset.xhtml?persistentId=doi%3A10.57745%2FYZCG1W&version=DRAFT. All R scripts used for analysis and figure generation are available on GitHub: https://github.com/benfield99/Romand-et-al.-2025-analysis-scripts/

---

**Figure S1.**
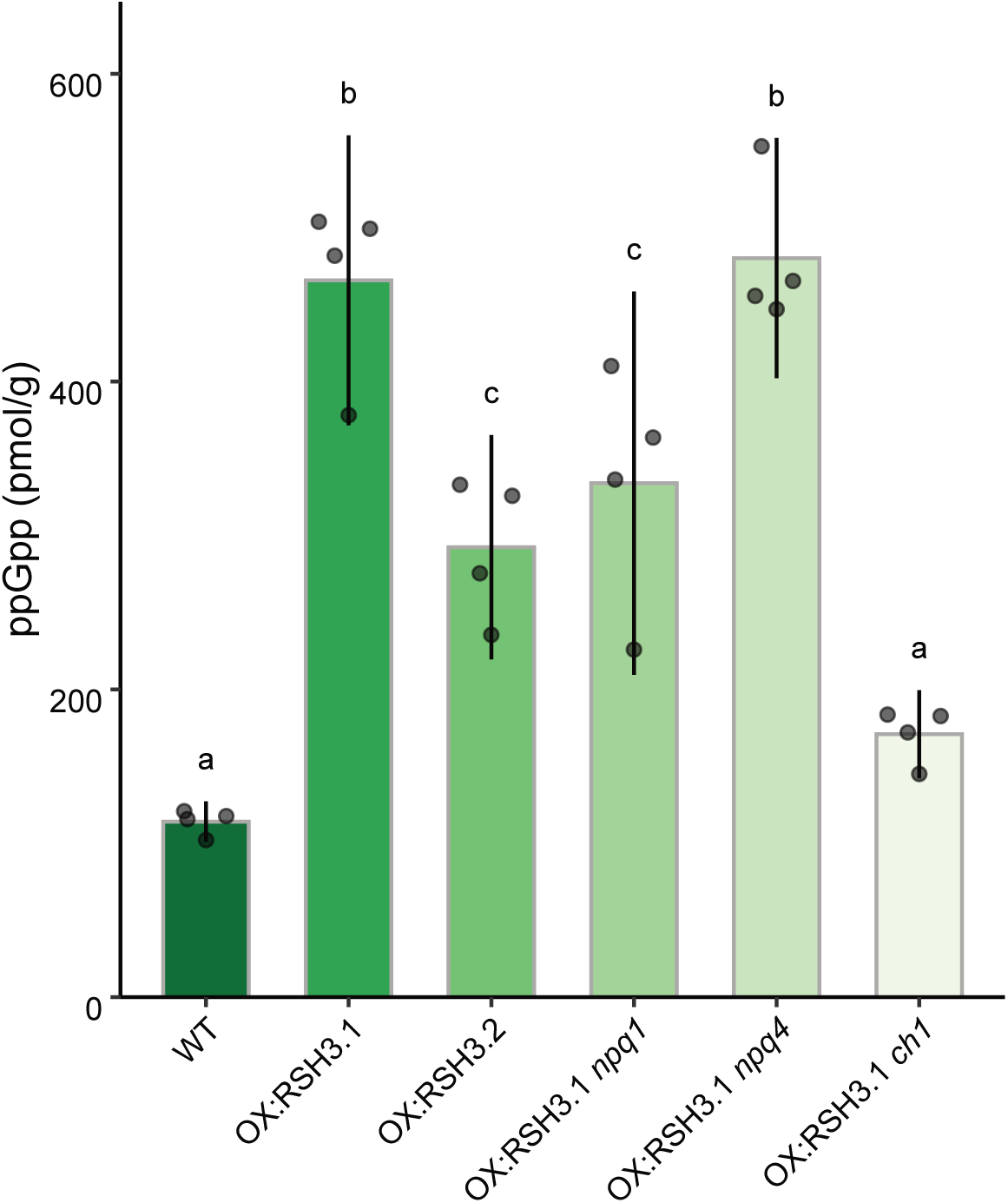
ppGpp levels in OX:RSH3 lines. ppGpp content in plants grown under standard conditions with growth light (GL) for 4 weeks. Graphs show mean ± 95% confidence interval, n = 4 biological replicates. Differences were tested using ANOVA and Tukey’s post-hoc test (see Materials and Methods). Letters indicate statistical groups. OX:RSH3.1 *ch1* did not maintain elevated ppGpp levels and was therefore not used to assess the effect of ppGpp in a *ch1* background. WT, wild type Col-0.

**Figure S2.**
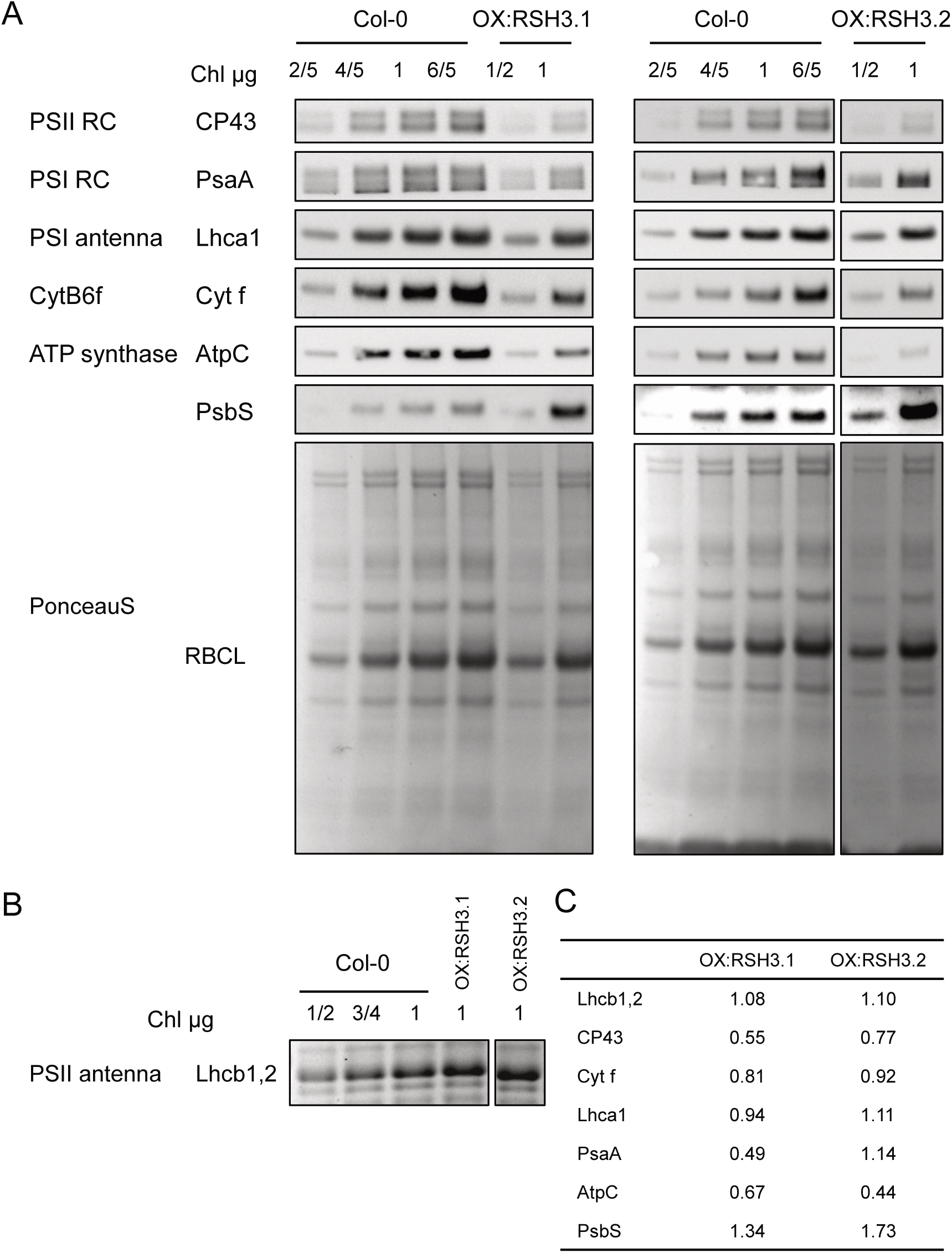
Photosynthetic protein levels in OX:RSH3 plants. (A) Immunoblots showing the abundance of the indicated proteins extracted from 4-week old plants. Total proteins were revealed using Ponceau stain (Ponceau S). (B) Lhcb1,2 abundance from Coomassie staining of thylakoid extracts in the indicated lines. (C) Quantification of relative protein amounts in OX:RSH3.1 and OX:RSH3.2 compared to the wild-type. Samples were normalised to total chlorophyll (Chl) content. Results are shown for two independent OX:RSH3 lines derived from a single experiment. PSII RC, PSII reaction centre; PSI RC, PSI reaction centre.

**Figure S3.**
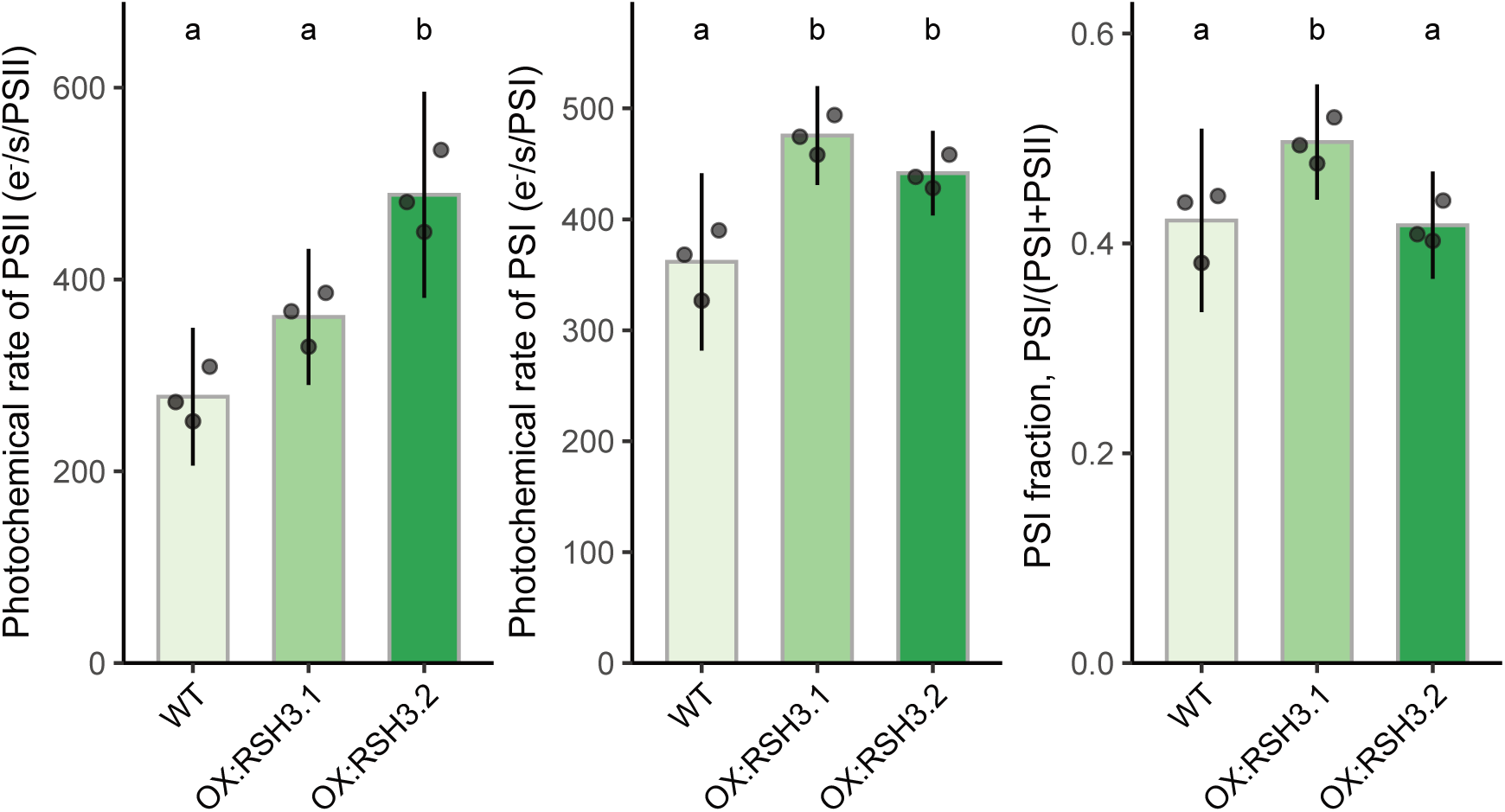
PS rates in OX:RSH3 lines. The photochemical rate of PSI and PSII and the PSI/(PSI+PSII) ratio were determined *in vivo* using electrochromic shift measurements. Means ± 95% confidence interval, n = 3 biological replicates). Differences were tested using ANOVA and Tukey’s post-hoc test. Letters indicate statistical groups.

**Figure S4:**
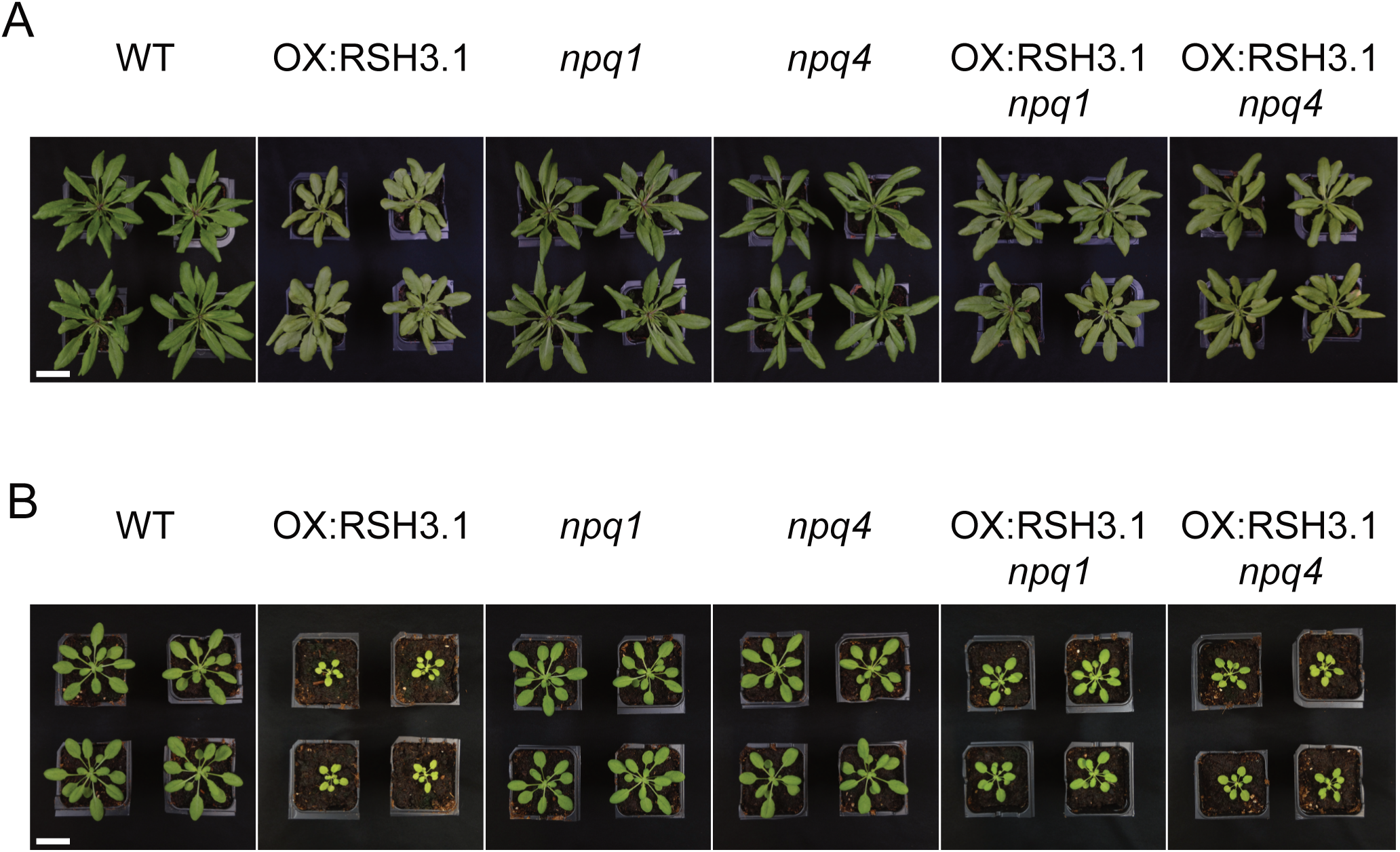
NPQ mutations partially suppress the growth defects of OX:RSH3.1. (A) Phenotypes of the different lines after 4 weeks of growth in long day conditions. (B) Phenotypes of the different lines after 4 weeks of growth in short day conditions. Scale bar, 3 cm.

**Figure S5.**
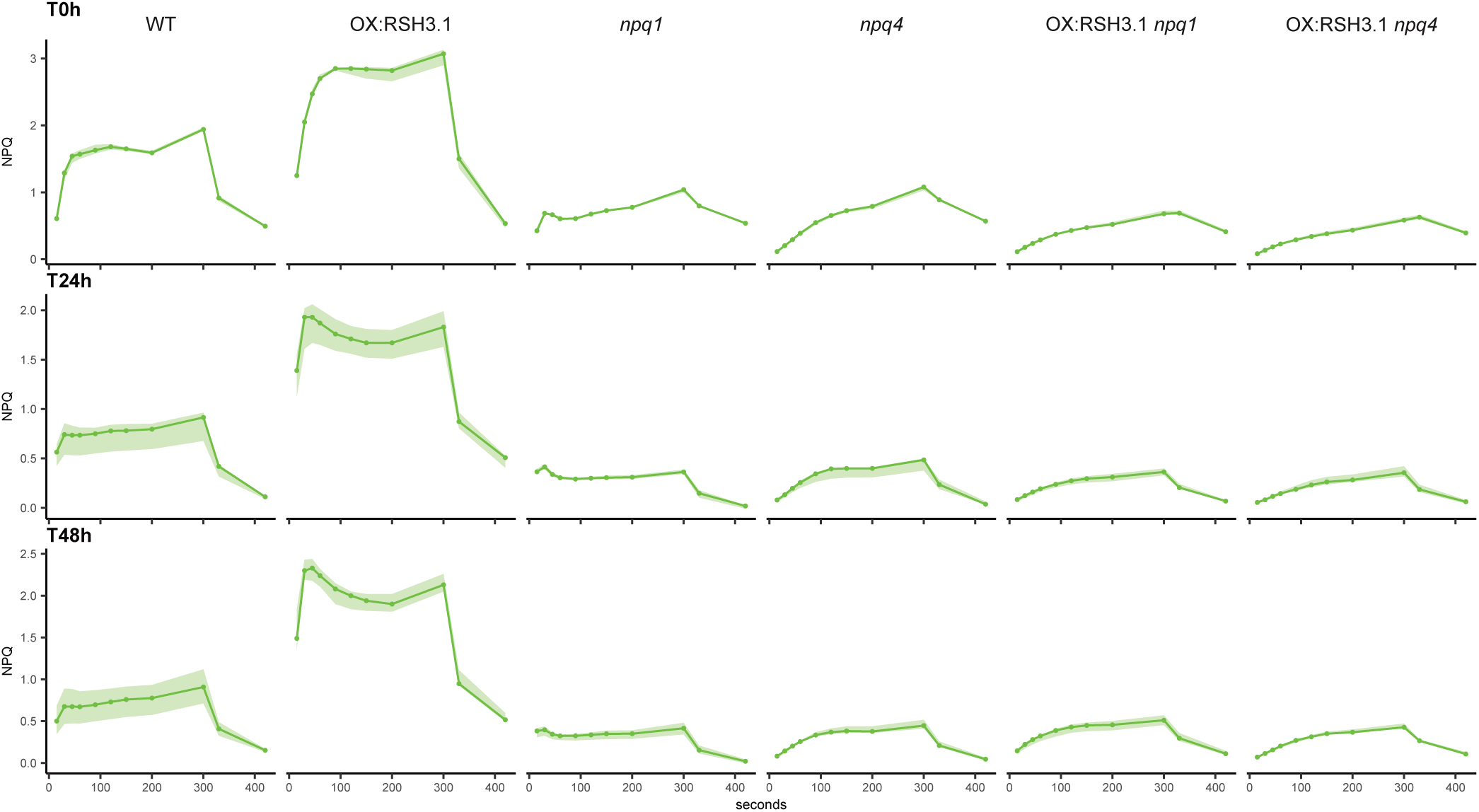
The effects of excess light exposure on NPQ. 4-week-old plants were exposed to excess light (EL; 1500 PPFD, air temperature 4–6 °C) and NPQ was measured by chlorophyll fluorescence imaging of individual plants after 24 h and 48 h of EL exposure. NPQ kinetics under an actinic light of 1200 PPFD. Curves represent median values ± 95% confidence interval, n = 4 biological replicates. Data shown in panels 1E, 2A, and S5 originate from the same experiment. PPFD, photosynthetic photon flux density (µmol photons m⁻² s⁻¹).

**Figure S6.**
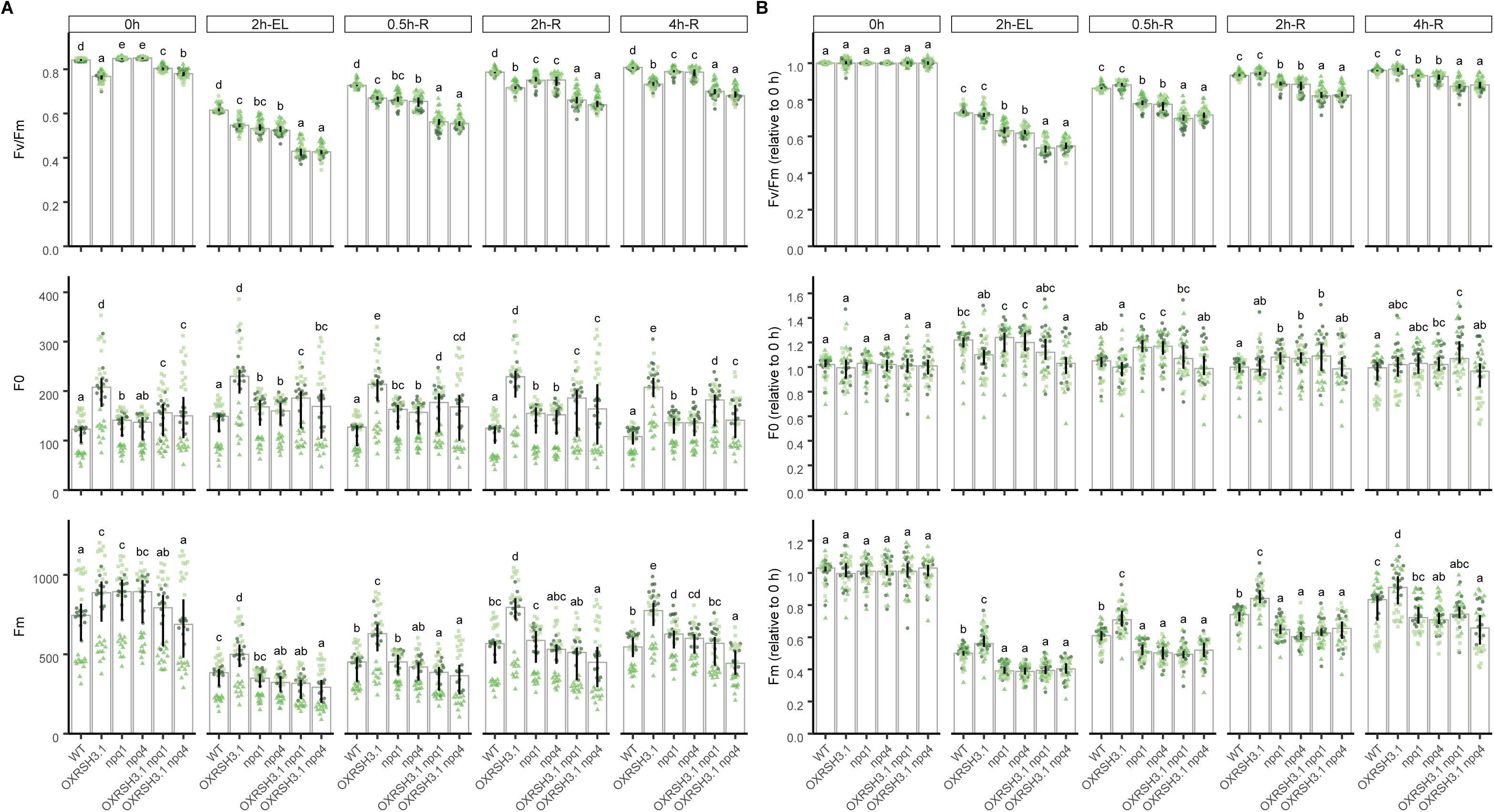
**Maximum quantum yield of PSII (Fv/Fm), minimal (F0) and maximal (Fm) fluorescence during short-term excess light exposure and recovery.** Data are from the experiment shown in Fig. 3c,d (see Fig. 3a for the protocol). (A) Absolute Fv/Fm (as in Fig. 3c), F0 and Fm. (B) Fv/Fm, F0 and Fm normalised to the mean value before EL exposure (0 h = 1). Graphs show medians ± 95% confidence intervals for three experimental replicates (circles, triangles and squares), each with 12 detached leaves per genotype. Letters indicate statistical groups between genotypes at each timepoint (aligned-ranks ANOVA with experimental replicate as a random effect, followed by pairwise comparisons, P < 0.05). Absolute F0 and Fm differ between replicates because of differences in instrument set-up, which does not affect Fv/Fm or the normalised values (B). Steady-state F0 and Fm cannot distinguish quenching from changes in antenna organisation or pigment content and should be interpreted with caution

**Figure S7.**
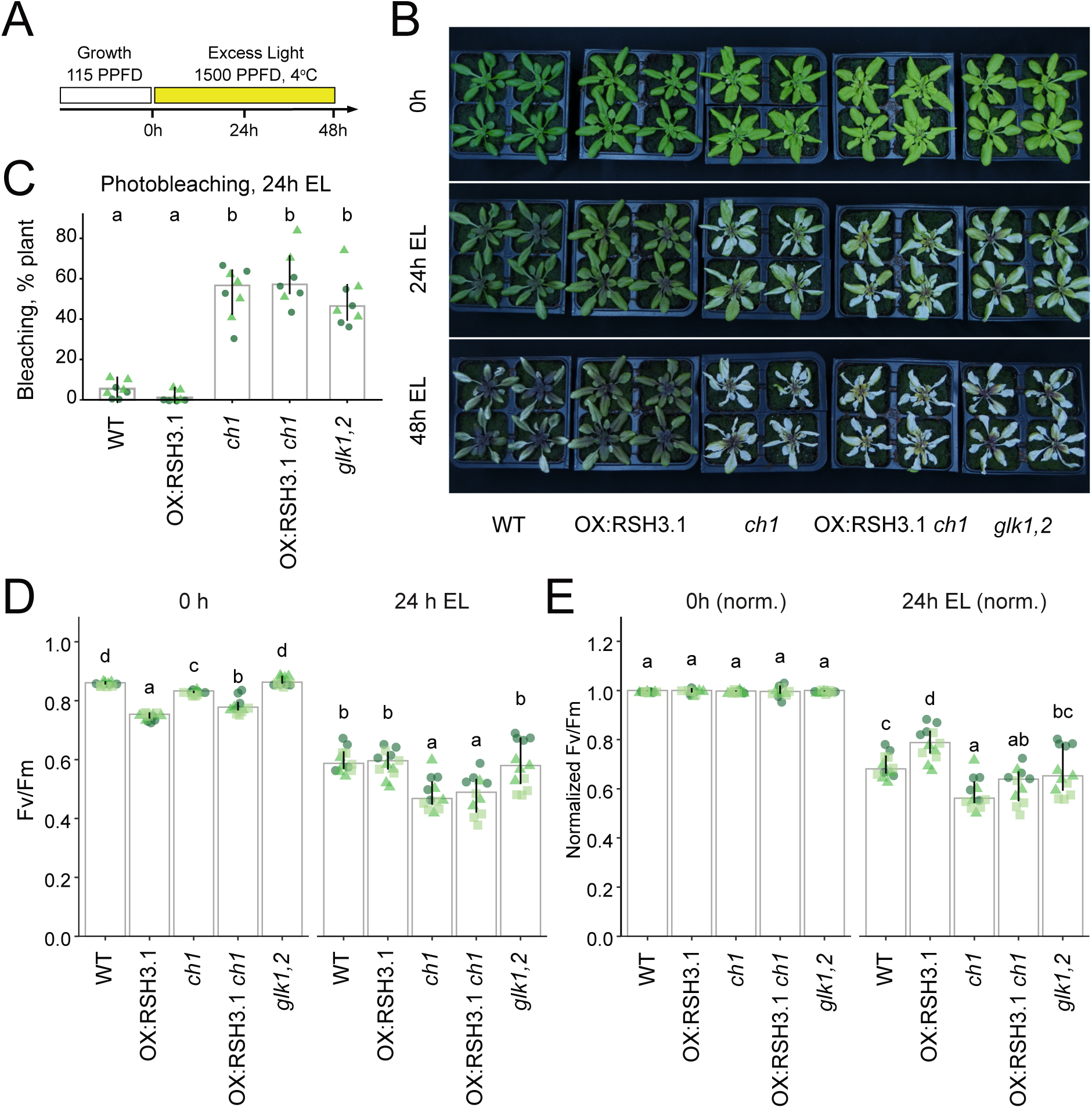
The low-chlorophyll mutants *ch1* and *glk1,2* are highly sensitive to excess light. (A) Experimental protocol. Plants grown to the same developmental stage were exposed to excess light (EL; PPFD 1500, air temperature 4–6 °C) for 48 h. (B) Plants were imaged to visualize photobleaching. (C) Quantification of photobleaching in the indicated lines after 24 h EL. (D) Maximum PSII yield was measured after 24 h EL. (E) Fv/Fm normalised to the median value before EL exposure (0 h = 1). Graph shows medians ± 95% confidence intervals for three experimental replicates (circles, triangles, and squares). Differences were tested using aligned-ranks ANOVA with post-hoc pairwise comparisons. Letters indicate statistical groups with comparisons between lines at each timepoint. OX:RSH3.1 *ch1* is shown for completeness. This line did not maintain elevated ppGpp levels (Fig. S1) and was therefore not used to assess the effect of ppGpp in a *ch1* background. PPFD, photosynthetic photon flux density (µmol photons m⁻² s⁻¹).

**Figure S8.**
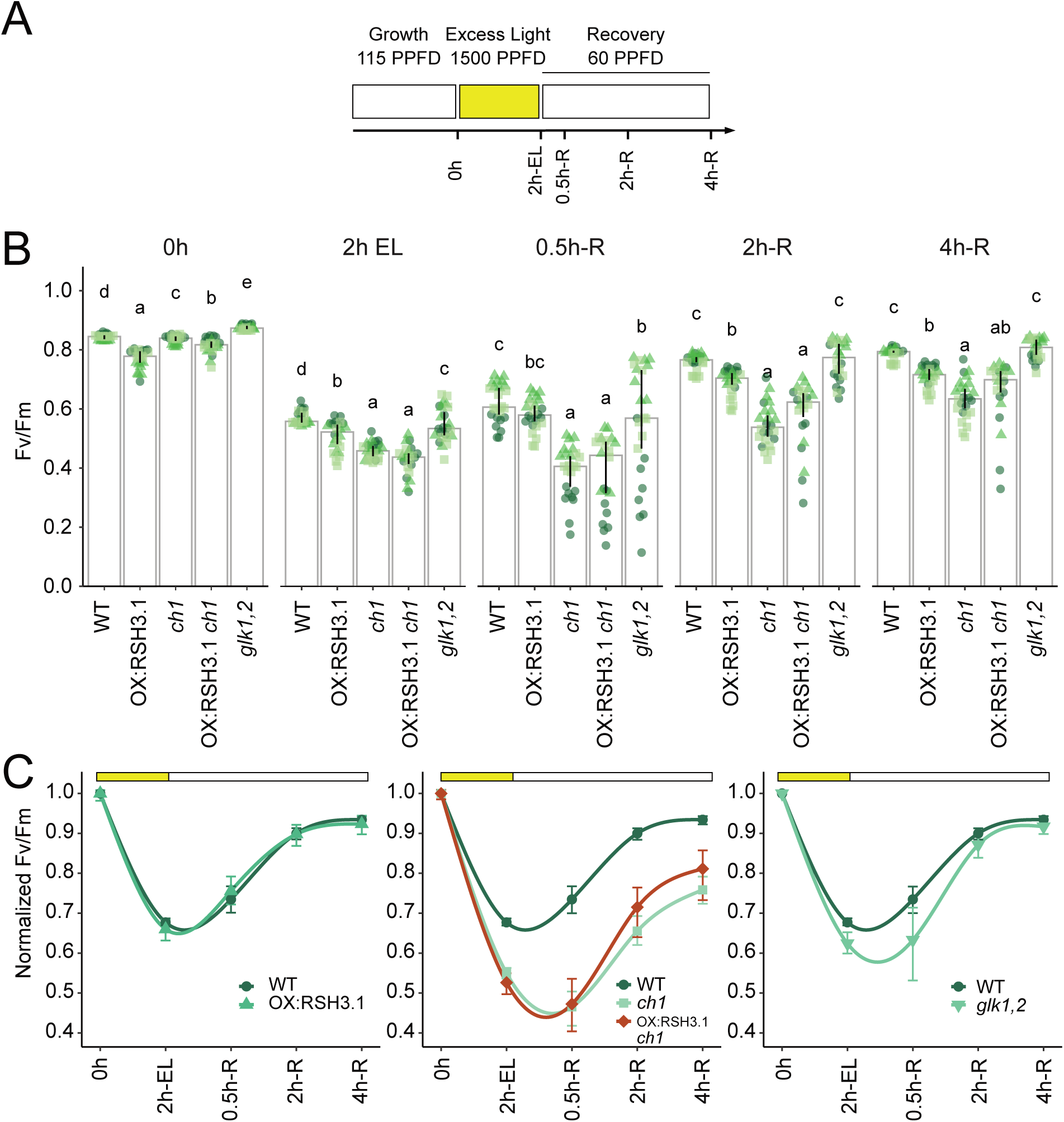
Photoinhibition and recovery of *ch1* and *glk1,2* during short-term excess light exposure. (A) Experimental protocol. Detached leaves of the indicated genotypes were exposed to excess light (EL; 1500 PPFD, air temperature 4–6 °C) for 2 h and then allowed to recover at 60 PPFD at 22°C. Timepoints: 0h, before EL; 2h-EL, after 2 h EL; 0.5h-R, 2h-R and 4h-R, after 0.5, 2 and 4 h of recovery. (B) Maximum PSII yield (Fv/Fm). Graphs show medians ± 95% confidence intervals for three experimental replicates (circles, triangles and squares), each with 12 detached leaves per genotype. Differences were tested using aligned-ranks ANOVA with post-hoc pairwise comparisons, letters indicate statistical groups between genotypes at each timepoint. (C) Fv/Fm normalised to the mean value before EL exposure (0 h = 1). Symbols show means ± 95% confidence intervals and lines are smoothed fits shown as a visual guide. The same wild-type data are shown in each panel to allow comparison with each genotype. OX:RSH3.1 *ch1* is shown for completeness; this line did not maintain elevated ppGpp levels (Fig. S1). PPFD, photosynthetic photon flux density (µmol photons m⁻² s⁻¹).

**Figure S9.**
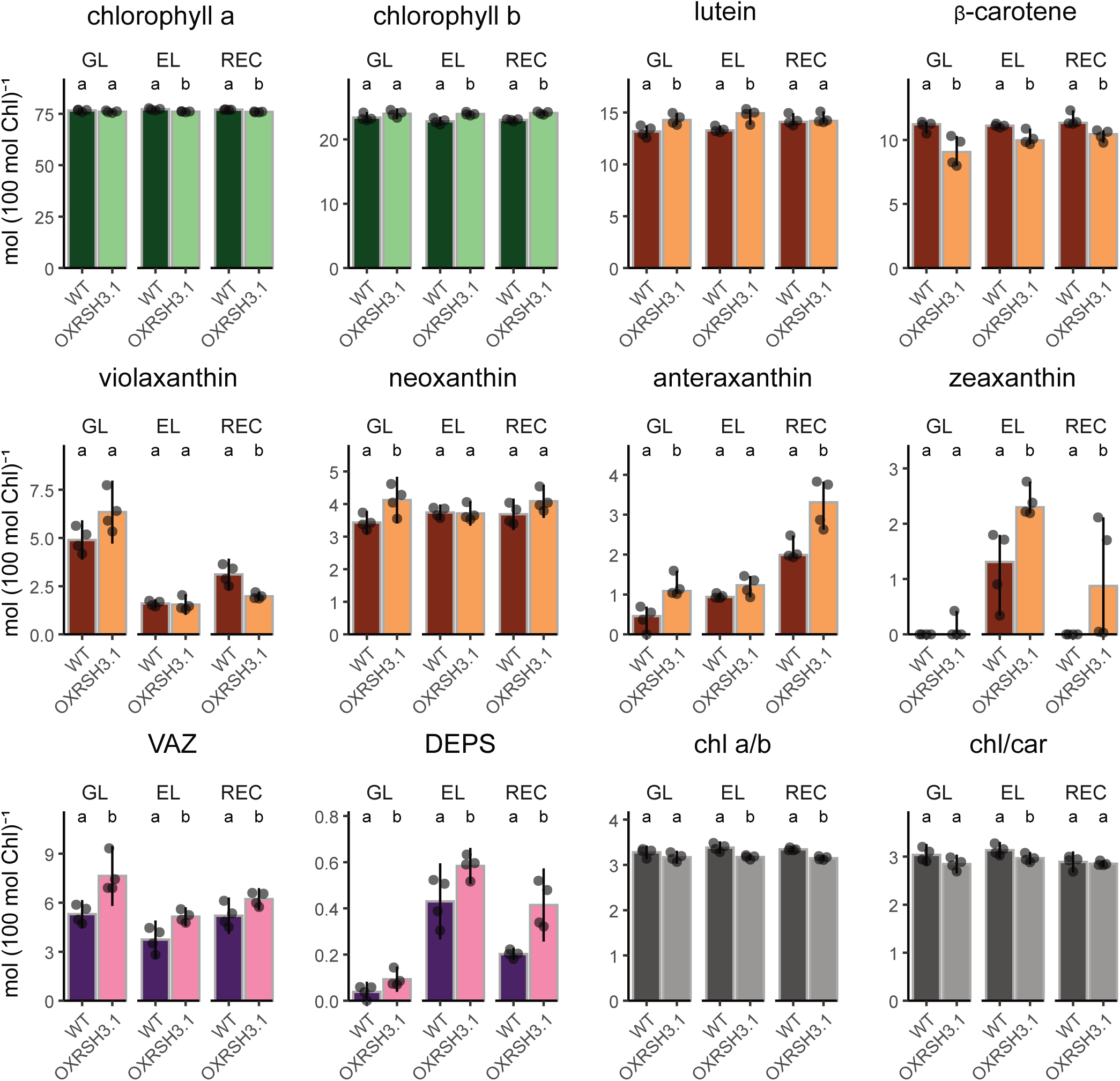
Pigment composition of wild type and OX:RSH3.1 plants before, during and after excess light. Four-week-old plants grown under growth light (GL) were exposed to excess light (EL; PPFD 1500, air temperature 4–6 °C) for 2 h and then allowed to recover under growth light for 1 h (REC). Pigments were extracted and analysed by HPLC. The first two rows show individual pigments. The last row shows the combined amount of violaxanthin, antheraxanthin and zeaxanthin (VAZ), the de-epoxidation state (DEPS = (Z + 0.5A)/VAZ)), the chlorophyll a/b ratio (chl a/b) and the ratio of total chlorophylls to total carotenoids (chl/car). Pigment contents (individual pigments and VAZ) are expressed in mol per 100 mol total chlorophyll (a+b); DEPS and pigment ratios are dimensionless. n = 4 biological replicates. Normally distributed data are shown as mean ± 95% confidence interval and were tested by ANOVA followed by Tukey’s post-hoc test. Non-normally distributed data (zeaxanthin) are shown as median ± bootstrap 95% confidence interval and were tested by the Kruskal-Wallis test. Letters indicate statistical groups between genotypes within each condition. PPFD, photosynthetic photon flux density (µmol photons m⁻² s⁻¹).

